# SnapKin: a snapshot deep learning ensemble for kinase-substrate prediction from phosphoproteomics data

**DOI:** 10.1101/2021.02.23.432610

**Authors:** Michael Lin, Di Xiao, Thomas A. Geddes, James G. Burchfield, Benjamin L. Parker, Sean J. Humphrey, Pengyi Yang

**Affiliations:** School of Mathematics and Statistics, The University of Sydney, NSW 2006, Australia; Computational Systems Biology Group, Children’s Medical Research Institute, The University of Sydney, Westmead, NSW 2145, Australia; Charles Perkins Centre, The University of Sydney, NSW 2006, Australia; School of Environmental and Life Sciences, The University of Sydney, NSW 2006, Australia; Department of Physiology, School of Biomedical Sciences, The University of Melbourne, Melbourne, VIC 3010, Australia

**Author notes:** To whom correspondence should be addressed: Pengyi Yang.

## Abstract

Mass spectrometry (MS)-based phosphoproteomics enables the quantification of proteome-wide phosphorylation in cells and tissues. A major challenge in MS-based phosphoproteomics lies in identifying the substrates of kinases, as currently only a small fraction of substrates identified can be confidently linked with a known kinase. By leveraging large-scale phosphoproteomics data, machine learning has become an increasingly popular approach for computationally predicting substrates of kinases. However, the small number of high-quality experimentally validated kinase substrates (true positive) and the high data noise in many phosphoproteomics datasets together impact the performance of existing approaches. Here, we aim to develop advanced kinase-substrate prediction methods to address these challenges. Using a collection of seven large phosphoproteomics datasets, including six published datasets and a new muscle differentiation dataset, and both traditional and deep learning models, we first demonstrate that a ‘pseudo-positive’ learning strategy for alleviating small sample size is effective at improving model predictive performance. We next show that a data re-sampling based ensemble learning strategy is useful for improving model stability while further enhancing prediction. Lastly, we introduce an ensemble deep learning model (‘SnapKin’) incorporating the above two learning strategies into a ‘snapshot’ ensemble learning algorithm. We demonstrate that the SnapKin model achieves overall the best performance in kinase-substrate prediction. Together, we propose SnapKin as a promising approach for predicting substrates of kinases from large-scale phosphoproteomics data. SnapKin is freely available at https://github.com/PYangLab/SnapKin.

## Introduction

Protein phosphorylation, one of the most pervasive cell signalling mechanisms, regulates a broad range of fundamental processes such as cell metabolism (*1*), differentiation (*2*) and the cell cycle (*3*), and its dysfunction has been linked to various diseases including cancers (*4*). Central to phosphorylation are the kinases that phosphorylate specific sites on their target substrate proteins. Together, kinases and their substrates establish the signalling networks of cells, governing all aspects of health and disease. Due to the significant time and resource cost on experimentally demonstrating the relationship between kinases and substrates, computational methods have been key workhorses for prioritising phosphorylation sites that are promising candidates prior to experimental verification. While many methods have been developed for predicting the cognate kinases of phosphosites, only a subset could perform kinase-specific predictions (*5*). Among the kinase-specific methods, most identify potential phosphorylation sites based on static information such as the amino acid sequences (*6*–*9*), protein structures (*10*), protein-protein interactions (*11*, *12*), or combinations thereof (*13*).

With recent major advances in mass spectrometry (MS)-based phosphoproteomics technologies, tens of thousands of phosphosites can now be quantified in a single experiment (*14*). These phosphoproteomics data provide a rich information resource that can be used for modelling the dynamics of each phosphorylation site in cells and tissues. Yet, very few computational methods utilise quantitative phosphoproteomics data for kinase-substrate prediction. These including CoPhosK, which uses co-phosphorylation patterns and interaction networks (*15*), DynaPho, which performs correlation analysis to identify kinase-substrate associations (*16*), and PUEL, an ensemble of support vector machine (SVM) models that predicts kinase substrates based on both kinase recognition motifs and phosphoproteomics dynamics (*17*). Nevertheless, the development of methods that extract information from phosphoproteomics data for kinase-substrate prediction is still in its infancy. Given the importance of phosphoproteomics in understanding the biology of cells, tissues, and complex diseases such as metabolic diseases and cancers (*18*), there is a growing need for advanced computational methodologies that utilise phosphoproteomics data to map these relationships.

Here, we aim to develop advanced machine learning models for kinase-substrate prediction by addressing several key challenges in learning from large-scale phosphoproteomics datasets. Specifically, to overcome the small number of experimentally validated kinase substrates, we introduce a ‘pseudo-positive’ learning strategy for increasing the size of training datasets during model building. To enhance model robustness and usage of training data, we implement a data re-sampling based ensemble learning strategy for classification models. To evaluate the models, we collect six published large phosphoproteomics datasets while also generating a new dataset from profiling muscle cell differentiation from mouse C2C12 myoblasts to myotubes. Using our collection of seven phosphoproteomics datasets and a panel of classification algorithms including both traditional and deep learning models, we first demonstrate the effectiveness of pseudo-positive and data re-sampling based ensemble learning strategies on improving model prediction and stability. In light of the utilities of these learning strategies on training deep learning models, we next propose an ensemble deep learning model (*19*), incorporating the pseudo-positive and data re-sampling strategies into a ‘snapshot’ ensemble learning algorithm (*20*). The resulting model, referred to as ‘SnapKin’, achieves the most competitive performance among all models evaluated for kinase-substrate prediction on all seven tested phosphoproteomics datasets. Lastly, we analyse the predictions from SnapKin on the new muscle cell differentiation phosphoproteomics dataset, highlighting the utility of phosphorylation profiles in complementing kinase recognition motifs in their substrate prediction. SnapKin is available as an R/Python package at https://github.com/PYangLab/SnapKin for kinase-substrate prediction from large-scale phosphoproteomics data.

## Results

Here we present the findings on using pseudo-positive and data re-sampling based ensemble learning strategies for improving model prediction and stability. Classification models included in the evaluation are naive Bayes (NB), logistic regression (LR); support vector machine (SVM), random forest (RF), XGBoost (XG), and densely connected neural network (DNN). We also benchmark the performance of the proposed SnapKin model, whereby the above two learning strategies are incorporated in a snapshot-based ensemble deep learning model. Lastly, we analyse the predictions from SnapKin on the muscle cell differentiation phosphoproteomics dataset, providing literature support for putative candidates uncovered by this computational model.

### Pseudo-positive strategy improves model prediction

A key limitation towards using supervised learning models for kinase-substrate prediction is the lack of high-quality positive training examples, owing to the small number of experimentally validated substrates for the majority of known kinases (*18*). While the number of phosphosites quantified in a phosphoproteomics experiment is often significantly larger than the number of substrates for which a kinase has been reported, only a subset may be used as negative training examples (through random subsampling in this study; see Materials and Methods). This is because many classification models are sensitive to class imbalance, where the binary classification of either positive or negative examples greatly outnumbers the other class (*21*). Since the substrates often show similar patterns of changes in phosphorylation upon the perturbation of their responsive kinases (e.g. stimulation, inhibition, differentiation) (*22*), we introduce a simple strategy in which for each kinase we select all pairs of its known substrates in the training dataset and average their phosphorylation profile to create additional positive training examples that we call ‘pseudo-positive’ examples (see Materials and Methods). The utility of these pseudo-positive examples can be assessed by evaluating the prediction performance of models on test datasets using cross-validation. Fig. 1 summarises the prediction performance of each model with and without the use of pseudo-positive examples. With the exception of the NB classifier, we found that the use of pseudo-positive examples resulted in significantly improved model performance in terms of area under the precision-recall curve (PR-AUC). These results demonstrate that the pseudo-positive strategy is effective for improving prediction across a range of classification models.

**Fig. 1.**
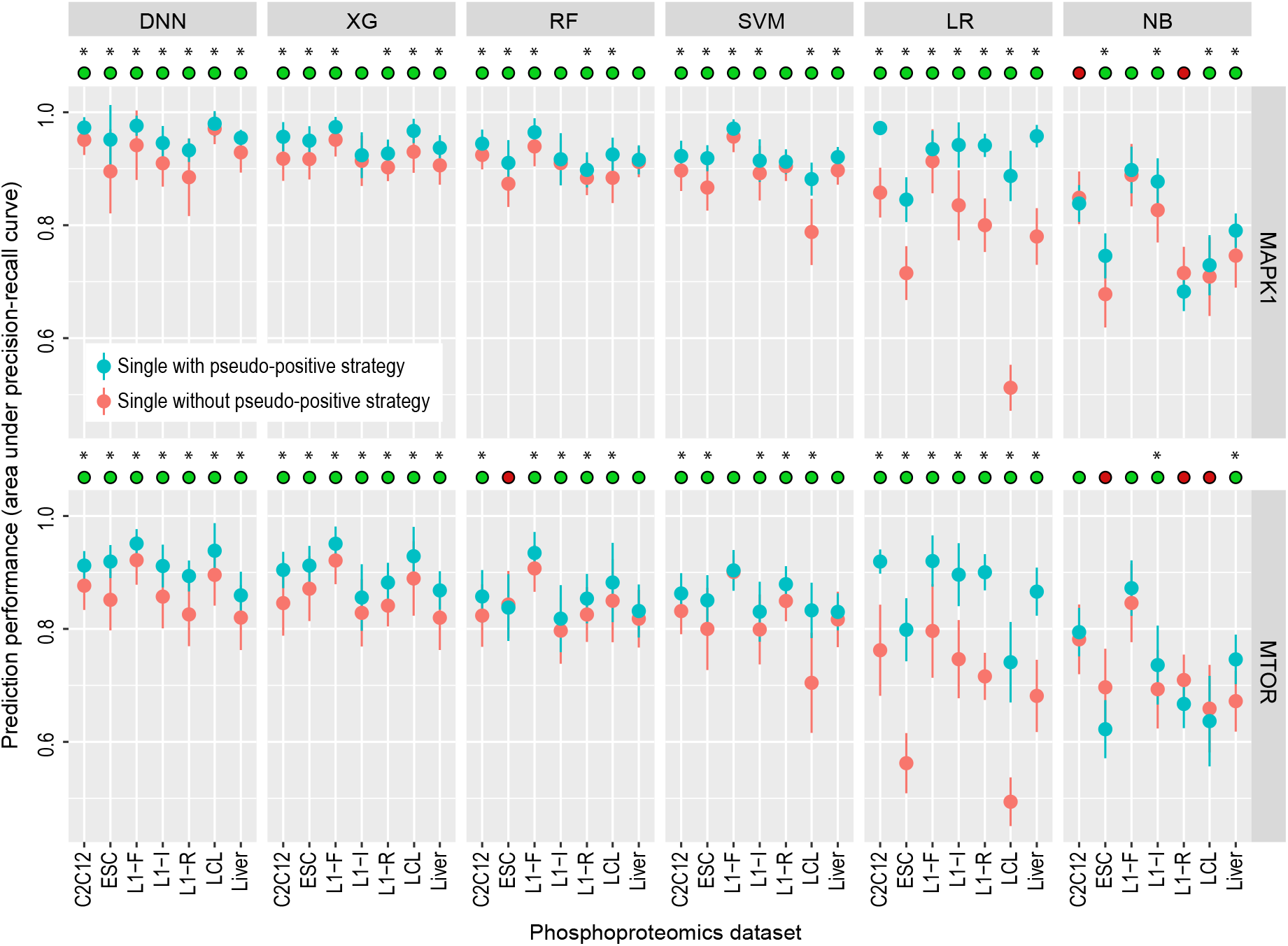
Prediction performance assessment of models with and without using the pseudo-positive learning strategy across the seven phosphoproteomics datasets. Solid dots represent the mean performance of each model from a 5-fold cross-validation and error bars represent standard deviation from 50 repeated trials of the 5-fold cross-validation. The green circles on top of each panel denote the cases when using the pseudo-positive strategy improves model performance and the red circles denote the opposite. * denotes *p*<0.05 using a one-sided Wilcoxon Rank Sum test.

### Data re-sampling based ensemble improves model prediction and stability

We propose a data re-sampling based ensemble learning strategy which involves generating multiple training datasets and consequently fitting multiple independent models in order to determine a collective prediction (*23*). In our kinase-substrate prediction setting, the motivation for the data re-sampling based ensemble learning stems from the need to utilise more of the negative training examples, given the large number of phosphosites quantified in the phosphoproteomics experiments, and the assumption that the majority of these are not substrates of a given kinase. For each given kinase, this is achieved by random subsampling from the phosphosites that are not annotated as its substrates and combining each sample with the positive examples to form multiple balanced training datasets from which multiple models are trained and combined to provide an ensemble prediction (see Materials and Methods for details). We compared the performance of models trained with and without using the data re-sampling based ensemble learning strategy and found in most cases a significant improvement in prediction is achieved when the model is trained using the ensemble learning strategy (Fig. 2). Another key advantage of ensemble learning is its robustness to data noise which can lead to more stable and reproducible predictions (*19*). Indeed, by comparing the variability in model prediction from the 50 repeated runs of the 5-fold cross-validation (see Materials and Methods), we observed a reduction of variance in most cases across the six models when the data re-sampling based ensemble strategy is used (Fig. 3A).

**Fig. 2.**
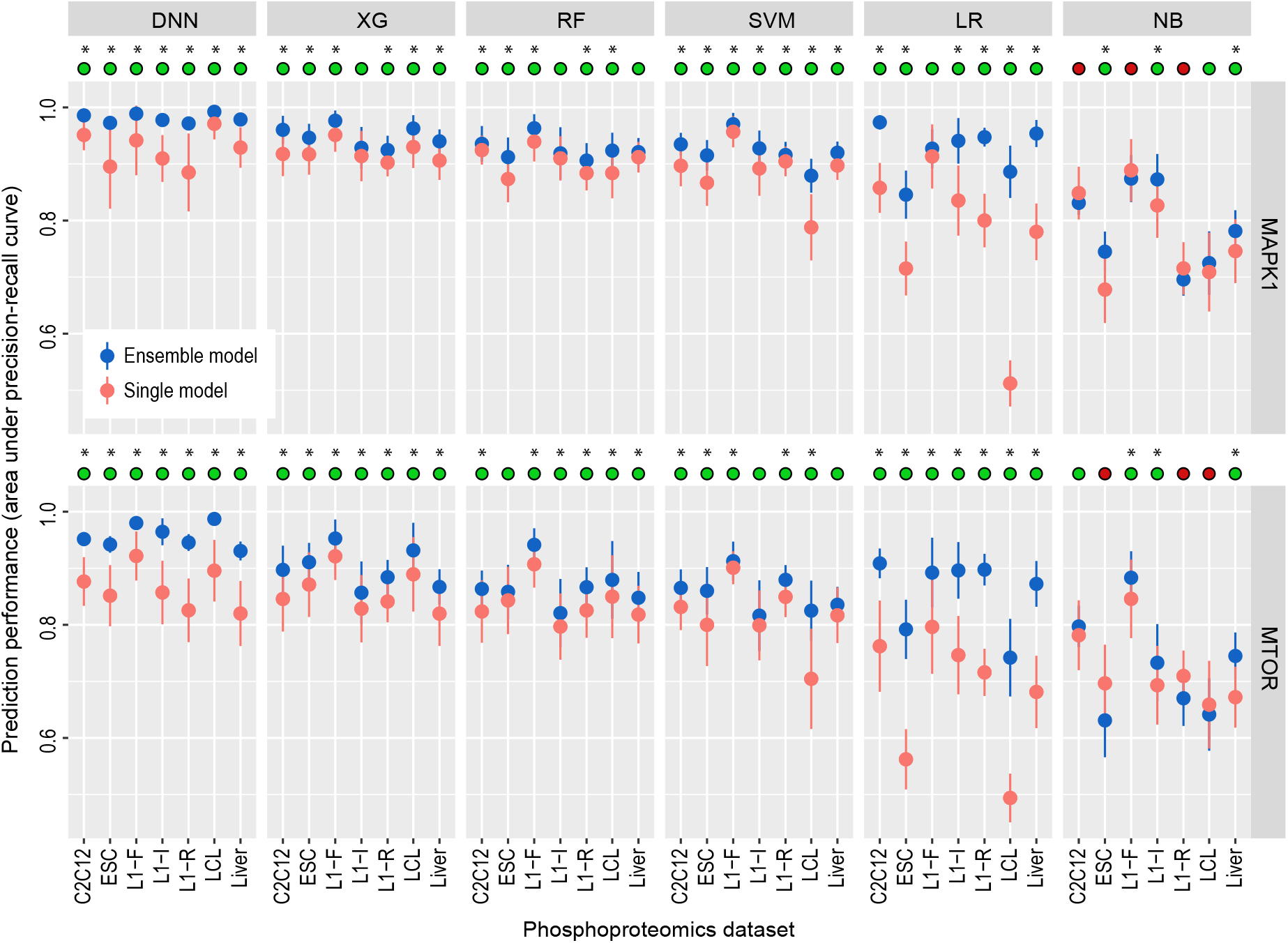
Prediction performance assessment of models with and without a data re-sampling ensemble learning strategy across the seven phosphoproteomics datasets. Solid dots represent the mean performance of each model from a 5-fold cross-validation and error bars represent standard deviation from 50 repeated trials of the 5-fold cross-validation. The green circles on top of each panel denote the cases when using the ensemble strategy improves model performance and the red circles denote the opposite. * denotes *p*<0.05 using a one-sided Wilcoxon Rank Sum test.

**Fig. 3.**
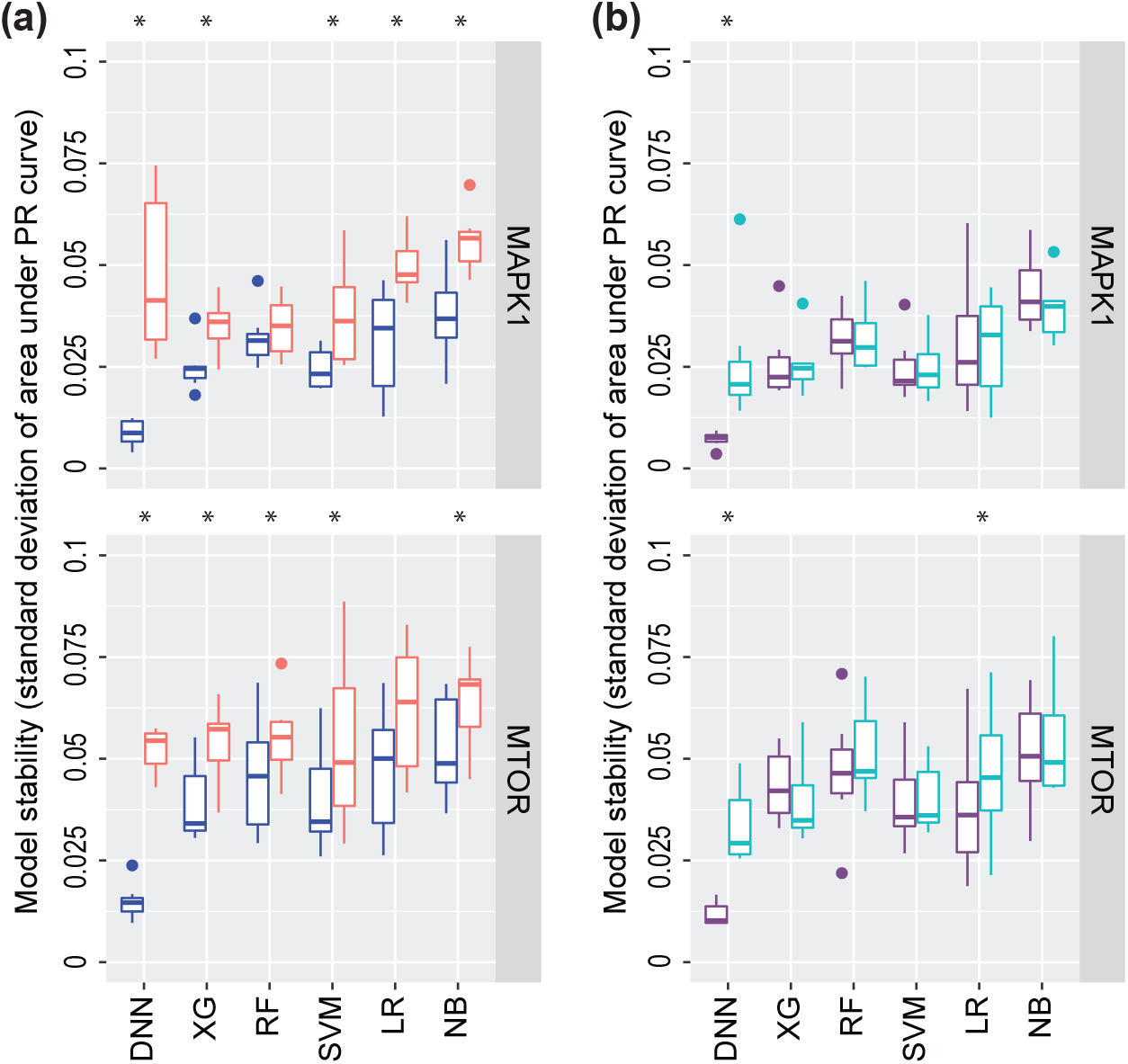
Assessment of model stability. (**A**) compares the stability of single and ensemble models trained using data re-sampling based ensemble strategy. (**B**) compares the stability of single and ensemble models trained in conjunction with the pseudo-positive strategy. Stability is quantified as the standard deviation of areas under the PR curves from the 50 repeated runs of the 5-fold cross-validation and each box contains the quantification from the seven phosphoproteomics datasets. * denotes *p*<0.05 using a one-sided Wilcoxon Rank Sum test.

Furthermore, the data re-sampling based ensemble strategy can be used in conjunction with the pseudo-positive learning strategy and may further improve model performance. To this end, we compared the prediction performance and stability of models trained using pseudo-positive examples and with or without using ensemble learning strategy. While the traditional classification models show no “synergistic” improvement from using both learning strategies, we found additional improvement for the deep learning model of DNN on both the model stability (Fig. 3B) and prediction (Fig. 4). These results are in line with the higher model complexity/flexibility of DNNs compared to traditional models, which may allow them to benefit more from additional training data.

**Fig. 4.**
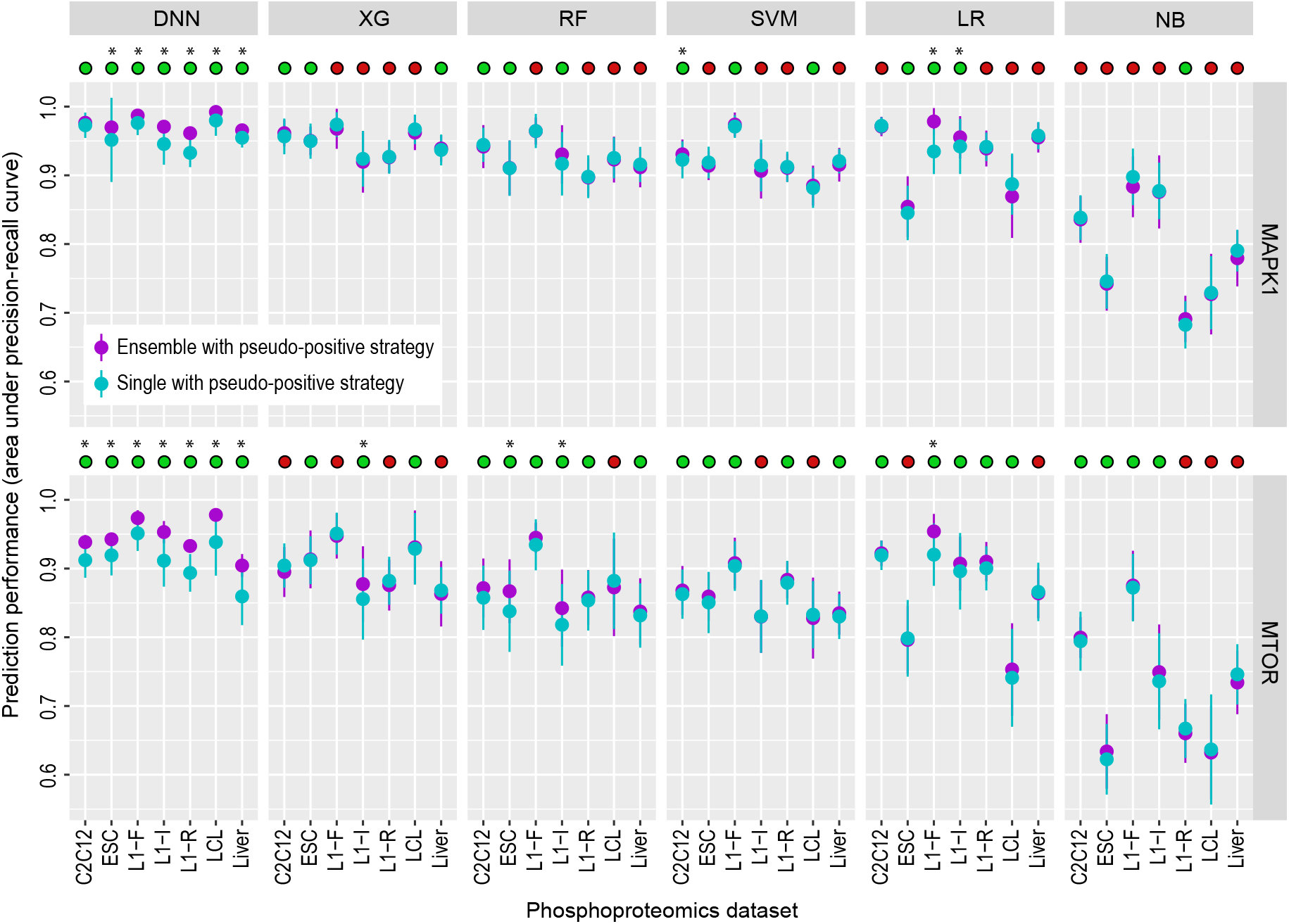
Prediction performance assessment of single and ensemble models trained in conjunction with the pseudo-positive strategy across the seven phosphoproteomics datasets. Solid dots represent the mean performance of each model and error bars represent standard deviation from 50 repeated runs of the 5-fold cross-validation. The green circles on top of each panel denote the cases when using the ensemble strategy improves model performance and the red circles denote the opposite. * denotes *p*<0.05 using a one-sided Wilcoxon Rank Sum test.

### Benchmark the performance of SnapKin

Our results from the above evaluation indicate that pseudo-positive and data re-sampling based ensemble learning strategies are effective towards improving model prediction and stability. They also demonstrate the competitive performance of the deep learning model (DNN) compared to traditional models, especially when used together with the two proposed learning strategies where additional performance gain is achieved mostly on DNN only (Figs. 3B and 4). In light of these findings, we further developed SnapKin, an ensemble deep learning approach wherein the pseudo-positive and data re-sampling strategies are incorporated into a snapshot ensemble model (see Materials and Methods). When compared to other models trained using pseudo-positive in conjunction with the ensemble learning, SnapKin shows the best overall prediction performance across all seven phosphoproteomics datasets and comparably small variability to the second best model (Fig. 5). Since in almost all cases, the second best method is DNN (trained using pseudo-positive and ensemble learning) which already has the smallest variability compared to traditional classification models (Fig. 3B), these results suggest that SnapKin achieves the best prediction performance without losing model stability compared to DNN. Given that in our implementation the DNN and SnapKin use the same network architecture, the performance improvement of SnapKin compared to DNN indicates that the snapshot ensemble, where a cyclic learning rate scheduler is utilised to perturb the network (*20*), brings further benefit on creating ensemble deep learning models in which various near-optimal models are extracted and combined in a single training process.

**Fig. 5.**
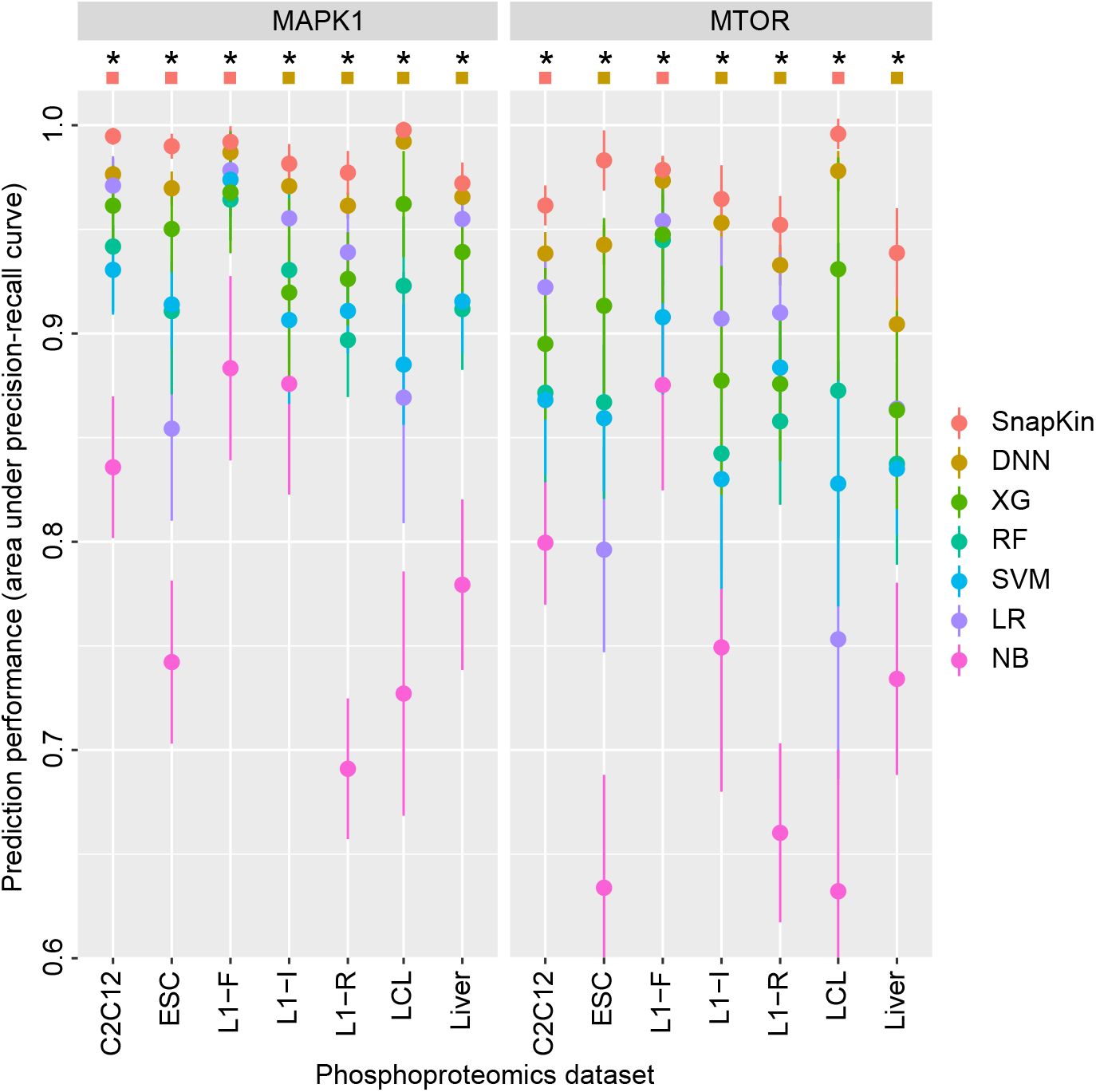
Prediction performance comparison of models across the seven phosphoproteomics datasets. Each model is trained using pseudo-positive and ensemble learning strategies, except SnapKin, which incorporates data re-sampling in each of the snapshot of the DNN (see Methods). Solid dots represent the mean performance and error bars represent standard deviation from 50 repeated trials of the 5-fold cross-validation. * denote *p*<0.05 comparing SnapKin with the second best method using a one-sided Wilcoxon Rank Sum test. Red squares denote when the standard deviation of SnapKin is smaller than the second best method whereas brown squares denote otherwise.

### SnapKin kinase-substrate predictions on muscle differentiation phosphoproteomics dataset

We next characterised the prediction results from SnapKin on the C2C12 differentiation phosphoproteomics dataset (Fig. 6A). We found that while most of the known MAPK1 and MTOR substrates have high prediction scores, the majority of the phosphosites in the dataset have close to zero prediction scores (Fig. 6B), consistent with the high selectivity of many kinases on their substrates (*24*). For putative MAPK1 and MTOR substrates predicted by SnapKin, the two groups show similar proline-directed consensus motifs (Fig. 6C), which are consistent with known MAPK1 and MTOR recognition motifs and common among many other kinases. Nevertheless, the phosphorylation profiles clearly distinguish the two groups with putative MAPK1 substrates showing acute phosphorylation increase at 30m time point and those of MTOR showing much slower response at day 5 (Fig. 6D). These results demonstrate that kinase recognition motifs alone may not be sufficient to identify kinase substrates and the phosphorylation profiles can be highly informative in distinguishing kinase-substrates that share similar motifs.

**Fig. 6.**
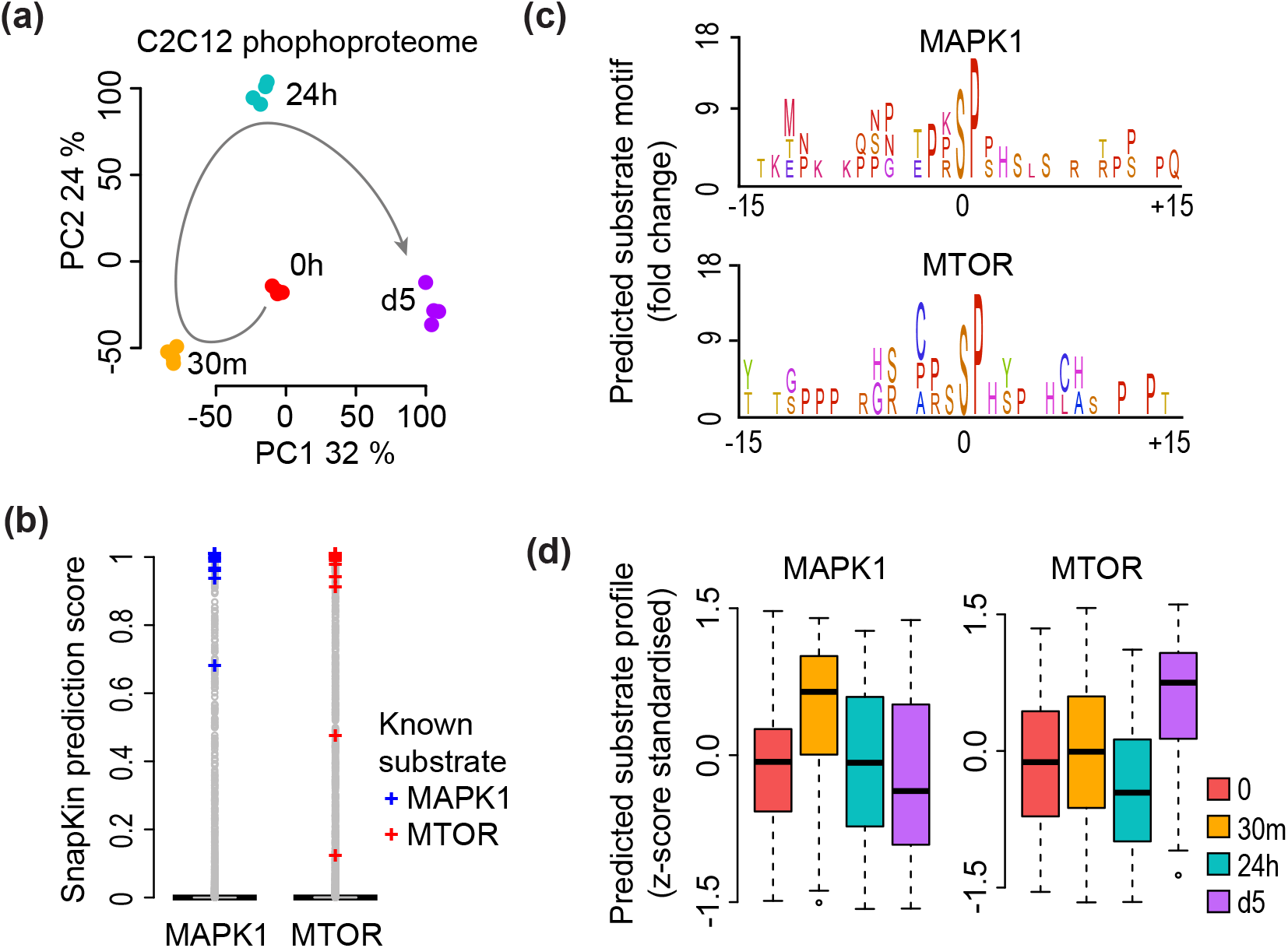
Muscle differentiation phosphoproteomics data analysis. (**A**) PCA visualising the phosphoproteomics data of C2C12 myoblast to myotube differentiation across four quantified time points from 0 (myoblasts), 30m, 24h, and day 5 (myotubes). (**B**) SnapKin prediction score on profiled phosphosites in C2C12 dataset. Known MAPK1 and MTOR substrates are highlighted in blue and red, respectively. (**C**) Consensus motif generated from SnapKin predicted MAPK1 and MTOR substrates. (**D**) Phosphorylation profiles from SnapKin predicted MAPK1 and MTOR substrates.

## Discussion

Global phosphoproteomics studies provide unprecedented opportunity to characterise signalling networks in health and diseases (*25*). While machine learning methods and especially deep learning algorithms can benefit from the abundant data generated from such studies, phosphoproteomic data-specific characteristics create various computational challenges limiting their direct application. One particular issue is the class imbalance caused by the small number of known kinase-substrate relationships because, compared to a small set of positive examples of a kinase, significantly more phosphosites can be used as negative examples for model training (*21*). Since most prediction models are sensitive to class imbalance, in this study, we have proposed various computational strategies to increase the size of the training dataset without introducing class imbalance. Nevertheless, other computational strategies such as cost-sensitive learning (*26*), which has been used for training classical neural networks (*27*), could be explored for developing ensemble deep learning models that alleviate the limit set by class imbalance, and may allow significantly more phosphosites to be included in training prediction models.

Typically, prediction models need to be trained using both positive and negative examples. For a kinase, although the positive examples can be found from known substrates such as those annotated in the PhosphoSitePlus database (*28*), the negative examples have to be defined independently as such information is often not available. Because only a relatively small number of phosphosites may be phosphorylated by each kinase owing to kinase-substrate selectivity (*24*), we treated the subsampled phosphosites that exclude the positive examples as negative examples, given that the chance of including unknown positive sites is small. While this assumption may have minimum effect on the comparison of model performance, including additional learning procedures that can take into account uncertainty in sampling negative examples may provide a more precise estimate of model accuracy (*29*) and will be explored in future work.

Related to the above, although the positive examples can be curated using known kinase substrates from an annotation database such as PhosphoSitePlus, there are various other databases (e.g. Phospho.ELM (*30*), PhosphoPOINT (*31*)) that can be used for such a purpose as well and the quality of the annotations may be dependent on the types of validation experiments and the biological systems in which they are validated. Developing methods that can take into consideration the type of evidence in kinase-substrate validation and the potential false positive examples in these data sources during model training will likely lead to further improvements in prediction accuracy.

While the experimental evaluation of kinase substrates remains time-consuming and labour-intensive, such data are nonetheless critical for validating predictions made from computational models. To this end, significant efforts have been made with the systematic mapping of kinase and their downstream substrates (*25*, *32*). Such experimental data resources will not only help validate putative kinase-substrate candidates from computational predictions but will also lead to improved predictive accuracy of computational models as the increasing number of experimentally validated kinase substrates will enable increasingly larger data repertoire to be curated for training computational models.

## Materials and Methods

This section provides the details for the generation of muscle differentiation phosphoproteomics dataset and the other public phosphoproteomics datasets used for model evaluation. It also details the implementation of classification models, learning strategies used for kinase-substrate prediction, and evaluation methods used for model assessment.

### C2C12 myoblast to myotube differentiation and sample preparation

C2C12 myoblasts were maintained in Dulbecco-minimum essential medium (DMEM) containing 25 mM glucose (Gibco) and 10% fetal bovine serum (Thermo Fisher Scientific) in a 5% CO2 incubator. C2C12 myoblasts were differentiated into myotubes over 5 days with 2% horse serum when myoblasts reached ~90-95% confluency. At the time points of 0, 30 minutes, 24 hours, and 5 days, cells were washed three times with ice-cold PBS and lysed in 4% sodium deoxycholate in 100mM Tris pH 8.5 and heated at 95 degrees before being snap frozen and stored at −20 degrees. After collection of all time points, cells were thawed, tip-probe sonicated and centrifuged at 16,000×g at 4 degrees to remove cellular debris. Protein was quantified with BCA (Thermo Fisher Scientific) and normalised to 240 ug followed by reduction with 10mM TCEP and alkylation with 40mM 2-chorloacetamide at 45 degrees for 5 min. Proteins were digested with 2.4 ug of sequencing grade trypsin (Sigma) and 2.4 ug of sequencing grade LysC (Wako) overnight at 37 degrees. Phosphopeptides were enriched using the EasyPhos protocol as previously described (*14*). Four biological replicates were generated for each time point.

### Phosphoproteomics analysis of myoblast to myotube differentiation

Phosphopeptides were separated on a Dionex 3500RS coupled to an Orbitrap Q Exactive HF-X (Thermo Scientific) operating in positive polarity mode. Peptides were separated using an in-house packed 75 μm×40 cm pulled column (1.9 μm particle size, C18AQ; Dr Maisch, Germany) with a gradient of 3-19% MeCN containing 0.1% FA over 20 min followed by 19-41% over 10 min at 350 nl/min at 55 degrees. MS1 scans were acquired from 350-1,400 m/z (60,000 resolution, 3e6 AGC, 50 ms injection time) followed by MS/MS data-dependent acquisition of the 10 most intense ions with HCD (15,000 resolution, 1e5 AGC, 50 ms injection time, 27% NCE, 1.6 m/z isolation width). Only multiply charged ions were selected for MS/MS with an apex trigger of 1-3 sec which were then excluded for 30 sec. Data was analysed with MaxQuant v1.6.12.0 (*33*) using all default parameters including 1% false discovery rates for peptide spectral matches and proteins. Methionine oxidation and Serine, Threonine and Tyrosine phosphorylation, and N-terminal protein acetylation were set as variable modifications while Cysteine carbamidomethylation was set as a fixed modification. Data was searched against the mouse UniProt database (August, 2019). The Phospho(STY) Sites table was processed in Perseus (*34*) to remove contaminants and reverse sequences followed by the ‘expand site’ function to obtain phosphosite-level quantification.

### Phosphoproteomics data processing

The muscle differentiation dataset generated from this study and six other public phosphoproteomics datasets generated from various cell types and tissues under experimental perturbations were used for model evaluation (table 1). Phoshoproteomics data from the C2C12 myoblast to myotube differentiation experiments were normalised and processed using PhosR package (*35*) for generating log2 fold change quantification at the four differentiation time points (Fig. 7A). For the other six public datasets, the processed phosphoproteomics quantification (in log2 fold change) were obtained from their respective studies. Since most supervised learning approaches rely on and generally perform better with more training data, we assessed the number of quantified kinase substrates in each dataset based on the known kinase-substrate annotation in the PhosphoSitePlus database (*28*). Fig. 7B shows that MAPK1 and MTOR are the two kinases with overall most quantified substrates (relative to other kinases in each dataset [min-max normalised]) across the seven datasets and were selected for subsequent prediction and evaluation experiments. For each phosphoproteomics dataset, we scored all phosphosites in the dataset based on MAPK1 and MTOR motifs using PhosR package and combined these motif scores with the phosphorylation dynamics (log2 fold change) using min-max scaling to form the input data for training each learning model.

**Table 1.** The phosphoproteomics datasets used in this study for kinase substrate prediction.

| **Dataset**<br>**(perturbation)** | **Abbreviation** | **# Phosphosite** | **# Feature** | **Accession** | **Publication** |
| --- | --- | --- | --- | --- | --- |
| C2C12<br>(differentiation) | C2C12 | 10495 | 18 | PXD023413 | This study |
| ESC<br>(differentiation) | ESC | 17866 | 50 | PXD010621 | (2) |
| L1 (FGF) | L1-F | 6864 | 14 | PXD003631 | (39) |
| L1 (Insulin) | L1-I | 12110 | 14 | NA | (40) |
| L1 (Redox) | L1-R | 17857 | 26 | PXD011525 | (41) |
| Liver cell lines<br>(Insulin) | LCL | 13330 | 26 | PXD001792 | (42) |
| Mouse liver<br>(Insulin) | Liver | 9687 | 93 | PXD001792 | (42) |

**Fig. 7.**
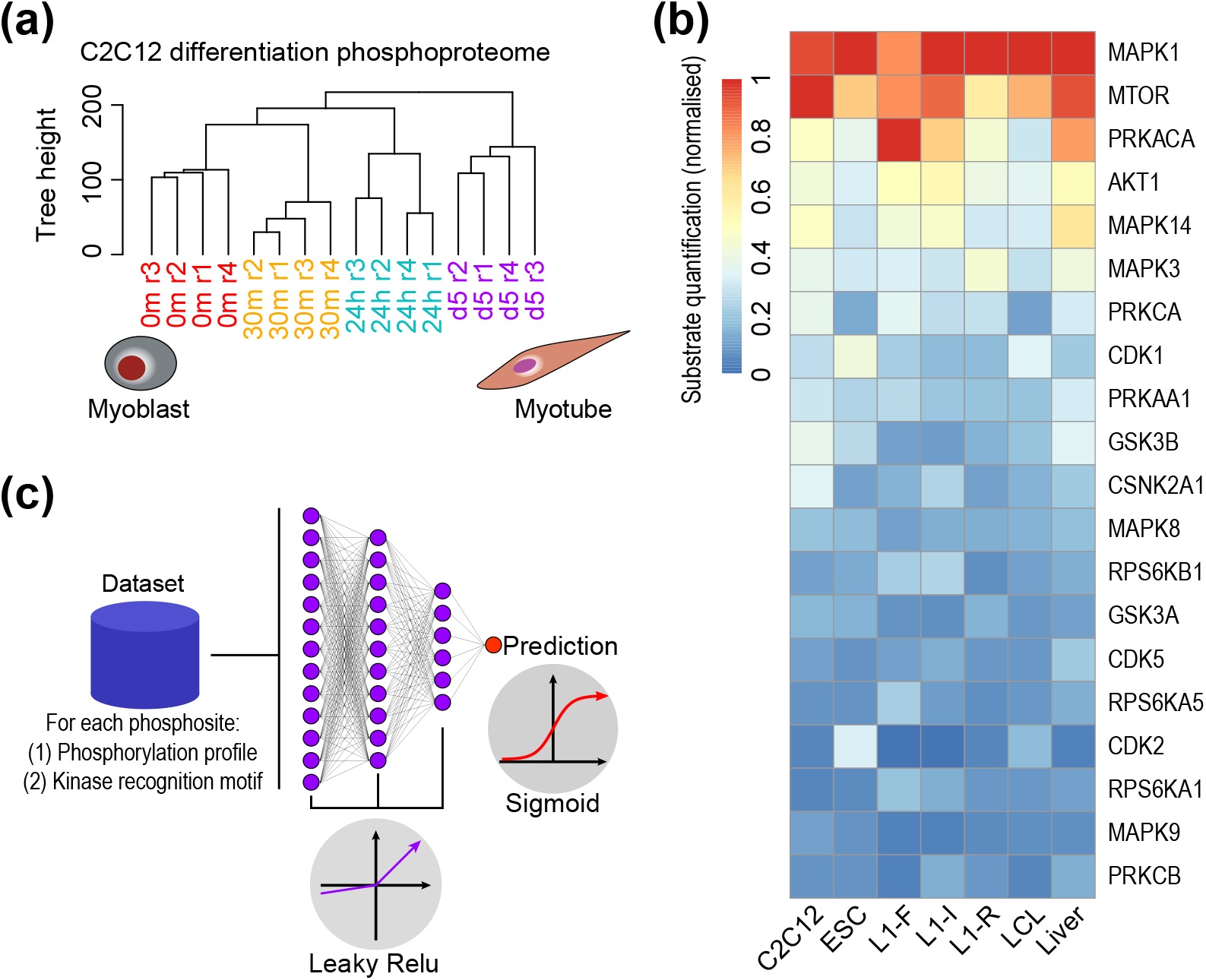
Phosphoproteomics data processing and deep learning neural network architecture. (**A**) Clustering of samples from the four time points (each in four biological replicates) profiled by MS-based phosphoproteomics during C2C12 differentiation. (**B**) Heatmap visualisation of the number of known substrates (based on PhosphoSitePlus database) quantified for a panel of kinases in each of the seven phosphoproteomics. (**C**) Schematic representation of the neural network architecture used in DNN and SnapKin.

### Classification models

We implemented a variety of classification models for testing their performance on kinase-substrate prediction. These include five traditional models and a deep learning model. For the traditional models, we implemented naive Bayes (NB), fitted using the the *discrim* R package; logistic regression (LR) using the *glm* R package; support vector machine (SVM) with a radial basis function using the *kernlab* R package; random forest (RF) with 500 trees using the *ranger* R package; and XGBoost (XG) with 1000 trees using the *xgboost* R package. For the deep learning model, we implemented a densely connected neural network (DNN) where we used fully connected neurons with hidden neurons activated by the ‘Leaky Relu’ function and the output neuron activated by a Sigmoid function (Fig. 7C). We found the hidden layers of three to be sufficient and determined the width of each layer using the following heuristic rules. We predefined a set of widths [2, 4, 8, 16, 32, 64, 128] and the first hidden layer of the DNN has a width equal to the largest value in the predefined width and less than or equal to the initial input features. Then it decreases by halving the width until the number of layers (i.e. 3) is reached. Other hyperparameters in our DNN include the ADAM optimiser (*36*), the binary cross-entropy loss function, epochs (i.e. 150), learning rates of 0.001, 0.01, or 0.1, and batch sizes of 32 or 64 obtained from a nested cross-validation of each fold.

### Pseudo-positive strategy

The small numbers of known substrates for most kinases introduce a major challenge in kinase-substrate prediction using phosphoproteomics data owing to the lack of positive training examples. This is further exacerbated by creating a balanced training dataset (*21*) since the small number of positive sites forces the subsampling of a small negative set. Hence, increasing the number of positive sites can lead to larger training dataset with both more positive and negative observations. Here, we propose the following steps for creating additional positive training examples:

1. Separate the phosphosites in the training dataset into the positive sites 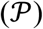 and the remaining sites that exclude the positive sites 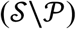;
2. For the *n*_*p*_ phosphosites in the positive set denoted by 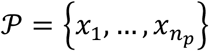, construct a list consisting of every unique pair of positive sites given by 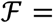 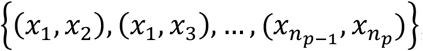;
3. For each pair in 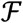, generate a pseudo-positive site *x*_*pseudo*_ = (*a* + *b*)/2, where 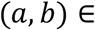 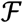. The pseudo-positive site can then be expressed as 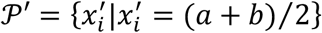;
4. The negative set 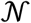 is then a subsample of 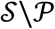 of the same size as the combined number of observations in 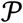 and 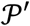. That is, 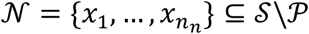 where 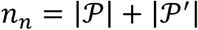;
5. The final training set is then the combined positive, pseudo-positive, and negative sites set 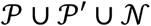.

This pseudo-positives strategy is able to generate *n*_*p*_(*n*_*p*_ − 1)/2 pseudo-positives meaning the subsequent adapted training dataset uses an additional *n*_*p*_(*n*_*p*_ − 1)/2 negative sites. This is particularly useful for supervised learning approaches which perform poorly with small sample sizes. Since substrates for a particular kinase typically exhibit similar temporal profiles (*22*), the pseudo-positive examples generated using this strategy make biological sense and have similar phosphorylation patterns to known phosphosites of a kinase, and hence can help improving the performance of the supervised learning approaches.

### Data re-sampling based ensemble strategy

Ensemble learning is an effective approach for dealing with small training data and data noise (*23*). The application of ensemble learning in kinase-substrate prediction therefore can further alleviate the issue of small number of training examples while also enhancing the robustness of the model and the stability of their prediction. To achieve this, we implement a data re-sampling procedure to generate multiple training datasets for training a collection of models using each supervised learning algorithm, and compute a final prediction score from their collective predictions through model averaging. This framework involves choosing the number of models within the ensemble denoted by *n*_*e*_ (set as 10 in this study) and is implemented in the following steps:

1. Separate the phosphosites in the training dataset into the positive sites 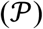 and other sites that are not the positive sites 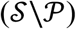;
2. Generate *n*_*e*_ separate training datasets denoted by 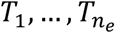, where each dataset 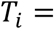 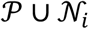 involves generating a new negative set 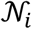 by repeated subsampling from 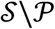, requiring 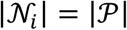;
3. For each training dataset, train a separate model *f*_*i*_(*x*|*T*_*i*_) for a total of *n*_*e*_ models;
4. To compute the prediction of a phosphosite *x*, compute the prediction for each model *f*_*i*_ and take the average of the prediction scores. Denote *F* to be the prediction from the ensemble model. The prediction is then defined by 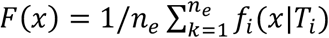.

This framework allows for an increased usage of 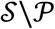 in training a model since 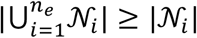 for each *i*. Additionally, by also including set 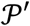 in the above pseudo-positive procedure for each training set, the ensemble procedure can be in conjunction with the pseudo-positive procedure where each 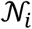 will have a size of 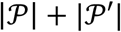.

### Implementation of SnapKin

The SnapKin model adopts the same architecture as in the above DNN but uses the stochastic gradient descent (SGD) and a learning rate scheduler (*20*) defined as the following:

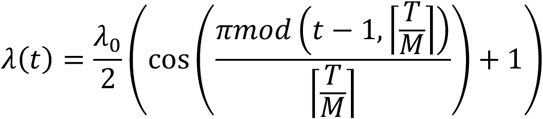

where *λ*_0_ was the initial learning rate (set as 0.01), *T* is the number of iterations (set as 1000), and *M* (set as 10 in this study to match the ensemble of DNNs) is the number of snapshots of the DNN. In addition, SnapKin adopts both pseudo-positive and data re-sampling learning strategies. Similar to the model ensemble strategy described above, a subsampling of the unannotated sites in a given dataset is performed to generate a training set 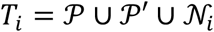 prior to training (*i* = 1) and after each snapshot is taken (*i* = 2 … *M*) and therefore enables better usage of data without introducing further computational time and model complexity, allowing our modification adhere to the ‘train 1, get M for free’ spirit of the original snapshot ensemble algorithm.

### Model evaluation

We applied a stratified *k*-fold cross-validation procedure for evaluating model performance. Specifically, we used *k*=5 in this study and repeated the cross-validation process 50 times to quantify the variability of model predictions. By stratifying each fold of the data, we ensure, for a given kinase, each fold maintains the ratio of positive and negative phosphosites in the original dataset. Each method was evaluated on each test fold of each phosphoproteomics dataset using the precision-recall (PR) curve defined by the four quantities: true positive (TP), true negative (TN), false positive (FP), and false negative (FN). PR curve is commonly used for comparing model performance especially when the dataset is highly imbalanced (*37*). It is a trade-off between *Precision*(*p*) = *TP*(*p*)/(*TP*(*p*) + *FP*(*p*)) and *Recall*(*p*) = *TP*(*p*)/(*TP*(*p*) + *FN*(*p*)), where *p* is the prediction threshold from each classifier. While the PR curves provide a threshold-based comparison of the models, we also used the areas under the PR curves as summaries and averaged them across all test folds in the cross-validation for quantifying overall performance of each model on each phosphoproteomics dataset. This allows us to easily compare models using statistical testing. Specifically, we used a one-sided Wilcoxon Rank Sum test with the hypotheses that (i) *H*_*a*1_: pseudo-positive strategy improves prediction of single models; (ii) *H*_*a*2_: ensemble learning improves prediction of single models; and (iii) *H*_*a*3_: ensemble learning in conjunction with pseudo-positive improves prediction on single model trained with pseudo-positive strategy. The areas under the PR curves from the 50 repeated runs of the 5-fold cross-validation were used as the primary statistics to compute the significance.

Finally, we used the standard deviation in the areas under the PR curves from the 50 runs of the 5-fold cross-validation to quantify the stability of the models. We then tested if the standard deviation from using ensemble learning is significantly smaller than single models across the seven phosphoproteomics datasets.

### Characterising SnapKin predictions on muscle differentiation phosphoproteomics dataset

Principal component analysis (PCA) was used to visualise the phosphoproteomes profiled from the four biologically replicates across the four differentiation time points. IceLogo (*38*) was used for generating consensus motifs from SnapKin predicted substrates (>0.8) for MAPK1 and MTOR, respectively. The SnapKin predicted substrates for MAPK1 and MTOR were also visualised for their temporal profiles using *z*-score standardised log2 fold change of phosphorylation compared to 0 time point.

## Acknowledgments

We thank the colleagues from the School of Mathematics and Statistics for their valuable feedback.

## Author contributions

PY conceptualised this work with input from SJH. ML and PY developed the methods with input from TAG. BLP performed myotube differentiation experiments and MS analysis. ML and DX performed the analysis with the supervision of PY and input from BLP and JGB. PY drafted the manuscript. All authors edited and approved the article.

